# Nfeature: A platform for computing features of nucleotide sequences

**DOI:** 10.1101/2021.12.14.472723

**Authors:** Megha Mathur, Sumeet Patiyal, Anjali Dhall, Shipra Jain, Ritu Tomer, Akanksha Arora, Gajendra P. S. Raghava

**Affiliations:** Department of Computational Biology, Indraprastha Institute of Information Technology, Okhla Phase 3, New Delhi-110020, India

**Author notes:** **Emails of authors**, Megha Mathur (MM), Sumeet Patiyal (SP), Anjali Dhall (AD), Shipra Jain (SP), Ritu Tomer (RT), Akanksha Arora (AA), Gajendra P. S. Raghava (GPSR). Contributed Equally (Megha Mathur, Sumeet Patiyal, Anjali Dhall, Shipra Jain). **Corresponding author**, Prof. G.P.S. Raghava, Head of Department, Department of Computational Biology, Indraprastha Institute of Information Technology, Okhla Phase 3, New Delhi-110020, India. E-mail address.

**Keywords:** Nucleotides, Composition-based Features, Software, Binary Profiles, Correlation-based Features

## Abstract

In the past few decades, public repositories on nucleotides have increased with exponential rates. This pose a major challenge to researchers to predict the structure and function of nucleotide sequences. In order to annotate function of nucleotide sequences it is important to compute features/attributes for predicting function of these sequences using machine learning techniques. In last two decades, several software/platforms have been developed to elicit a wide range of features for nucleotide sequences. In order to complement the existing methods, here we present a platform named Nfeature developed for computing wide range of features of DNA and RNA sequences. It comprises of three major modules namely Composition, Correlation, and Binary profiles. Composition module allow to compute different type of compositions that includes mono-/di-tri-nucleotide composition, reverse complement composition, pseudo composition. Correlation module allow to compute various type of correlations that includes auto-correlation, cross-correlation, pseudo-correlation. Similarly, binary profile is developed for computing binary profile based on nucleotides, mono-nucleotides, di-/tri-nucleotide properties. Nfeature also allow to compute entropy of sequences, repeats in sequences and distribution of nucleotides in sequences. In addition to compute feature in whole sequence, it also allows to compute features from part of sequence like split, start, end, and rest. In a nutshell, Nfeature amalgamates existing features as well as number of novel features like nucleotide repeat index, distance distribution, entropy, binary profile, and properties. This tool computes a total of 29217 and 14385 features for DNA and RNA sequence, respectively. In order to provide, a highly efficient and userfriendly tool, we have developed a standalone package and web-based platform (https://webs.iiitd.edu.in/raghava/nfeature).

## Introduction

In recent years, the amount of available biological sequence data has increased exponentially due to the significant increase in genome sequencing projects (Benson et al., 2013; Cochrane et al., 2016; Collins & Fink, 1995; Hood & Rowen, 2013; Leinonen et al., 2011; Pareek et al., 2011; Schatz, 2015; Stoesser et al., 2002). With an increasing data set, it is important to capture relevant information to solve biological questions (Brodland, 2015; Sabyasachi Dash, 2019). In literature, several studies reported the application of machine learning techniques in annotation of biological molecules (DNA/RNA) (Huang et al., 2021; Jonathan Schmidt, 2019; Mahmud et al., 2021; Xu & Jackson, 2019; Yang et al., 2020). It is important to compute features of nucleotide sequences, in order to implement machine learning techniques for developing models for annotating biological molecules (Abdurakhmonov, 2016; Jurtz et al., 2017; Usman). In past, various computational tools are reported (such as PseKNC (Chen et al., 2014), RepDNA (Liu et al., 2015), RepRNA (Liu et al., 2016), BioTriangle (Dong et al., 2016), PyBioMed (Dong et al., 2018), BioSeq-Analysis2.0 (Liu et al., 2019)) for computing various sequence-based features for a given DNA/RNA sequence. Despite several methods developed in the past, some important features have not been integrated into those platforms.

In order to supplement previous efforts, we have made a systematic attempt to developed a webserver platform “Nfeature” that integrates most of the features discovered in the past along with the incorporation of new features. In this study, we have introduced new features Nucleotide Repeat Index (NRI), Distance distribution of Nucleotides (DDN) and Entropy at sequence level as well as nucleotide level as a novel feature for DNA/RNA sequences. We have also incorporated Binary profile-based features for a given nucleotide sequence. These features are essential for motif predictions, factors/enhancers binding sites, etc. Using these modules, user can easily calculate the binary fingerprints of each nucleotide in a given sequence. Nfeature is a comprehensive platform to fetch all relevant information from a given nucleotide sequence in the form of vectors, which can be directly used for developing prediction models. A user-friendly web server and standalone package have been developed to facilitate users in computing features of nucleotide sequences (Figure 1).

**Figure 1:**
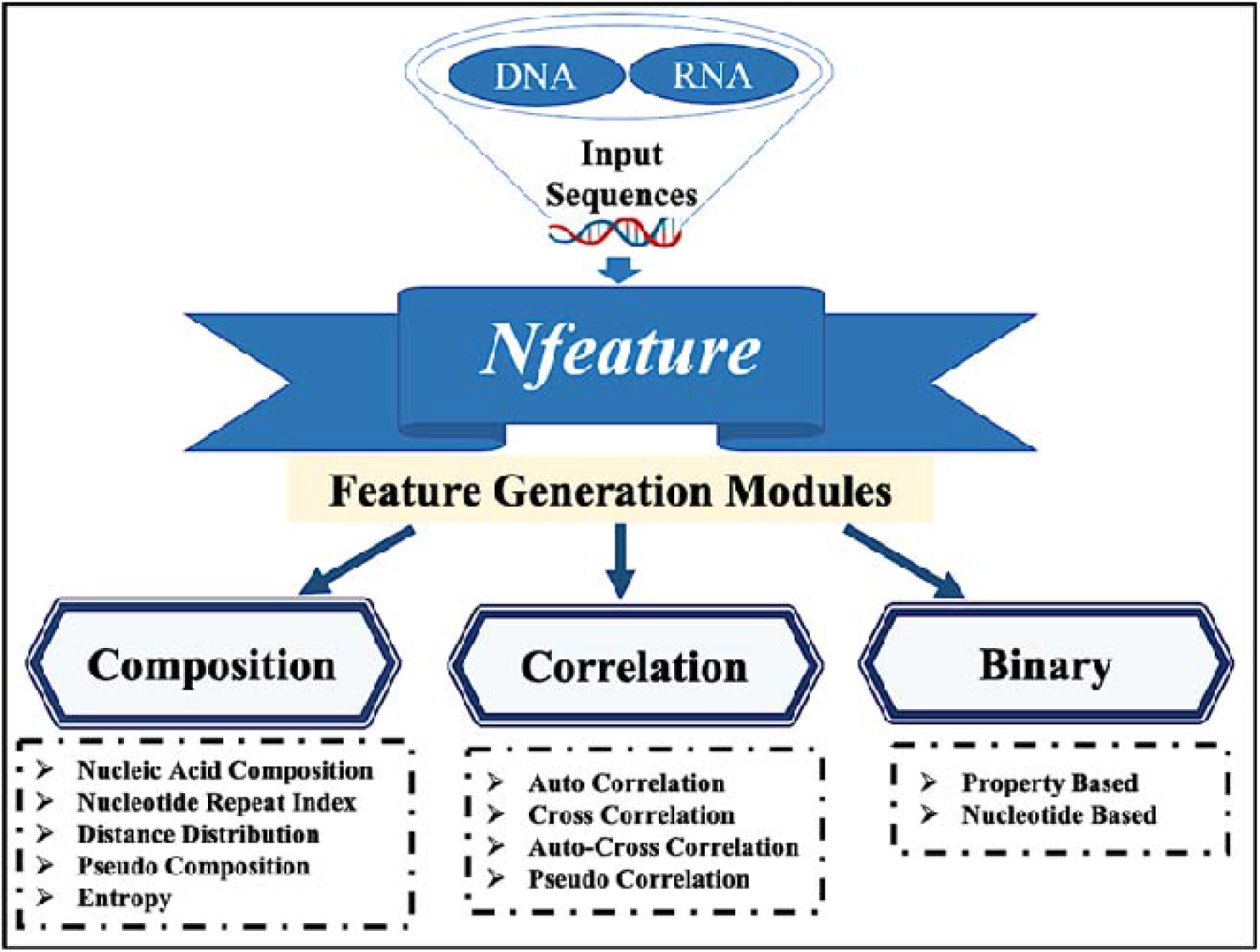
Schematic representation of major modules of Nfeature.

### Nfeature Overview

#### Composition/distance distribution-based features

This module aims to calculate nucleotide composition-based features in nucleotide sequences. It enables users to compute nucleic acid composition, distance distribution of nucleotides (DDN), nucleotide repeat index (NRI), pseudo composition and entropy of a sequence. We have incorporated most of the features used in previous studies. In addition, we integrate new features like entropy where we compute entropy at sequence and at nucleotide level. Several past studies have shown that nucleotide repeats have important biological functions. For example, repeated DNA residues are essential for the expression of unique coding sequence which further form nucleoprotein complexes. Whereas, in some cases these repeated sequences causes biological disorders like “GGGGCC” repeat sequences in the *C9orf72* gene causes neurodegenerative disease (Chanou & Hamperl, 2021; De Bustos et al., 2016; Handy et al., 2011; Malik et al., 2021; Miret et al., 1997; Rajewska et al., 2012; Shapiro & von Sternberg, 2005) (Abugable et al., 2019). Additionally, Pfeature, a computational tool of feature extraction also incorporates residue repeat and distance distribution feature generation methods for the protein sequences (Akshara Pande, 2019). Hitherto, no method capture such information of nucleotides from nucleotide sequences. In this study, first time we used NRI to calculate the repeating nucleotide information of DNA/RNA sequence. It measures the number of continuous runs of a nucleotide in a biological sequence. To calculate the distance distribution of nucleotide residues, a module is used for computing distance distribution information for a nucleotide sequence. Residue repeat and distance distribution of nucleotide sequence can be calculated by using the given formula:

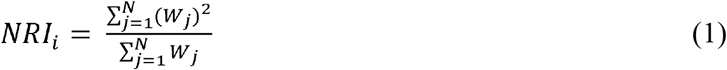

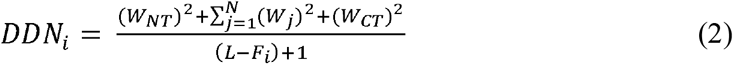

Here, NRI_i_ and DDN_i_ stands for nucleotide repeat index and distance distribution of nucleotide type i, where N stands for the maximum number of occurrences, W_j_ stands for the number of repeats in occurrence “j” for nucleotide type “i”, W_NT_- nucleotide distance from N-terminal; W_j_- Inter-distance between nucleotides “i”; W_CT_- Nucleotide distance from C-terminal, L- Total length of nucleic acid sequence; F_i_ – Frequency of nucleotide type “i”.

Shannon entropy plays a significant role in the field of information theory. Recently several studies have shown the importance of entropy in DNA sequences for example, to investigate exons and introns in the DNA sequences (Li et al., 2019), to identify DNA sequence diversity between different alleles within one individual (Sherwin, 2010). To the best of our knowledge there is no method incorporating entropy-based features. So, to get the Shannon entropy information of DNA and RNA sequences, we first time introduce an entropy-based module which computes entropy of DNA/RNA sequences at residue as well as sequence level. Sequence and residue level entropy is calculated using following equations 3, and 4.

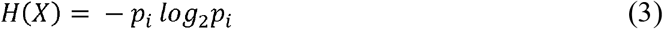

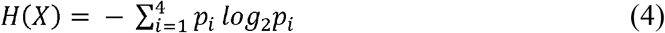

Here “i” is the nucleotide in the sequence and X is any nucleotide sequence, and p_i_ is the probability of a given nucleotide in the sequence.

#### Correlation based features

In this module, correlation-based features of DNA and RNA sequences are calculated. Correlation is defined as a relation between properties/features i.e. if a feature variable is related to its own then it is defined as auto-correlation and if there exists some correlation between two features/variables then it is known as cross-correlation. Correlation based features basically convert the different length DNA and RNA sequences into fixed length vectors, so that machine learning techniques can be applied to the extracted features. These descriptors identify features based on nucleotide properties along the sequence.

The correlation module of Nfeature calculates the diverse range of features for DNA and RNA sequences, including autocorrelation, cross-correlation, auto-cross correlation, and pseudo correlation. The autocorrelation module further subdivided into seven categories based on the properties and correlation types, such as dinucleotide based autocorrelation (DAC), trinucleotide-based autocorrelation (TAC), dinucleotide based Moran autocorrelation (DMAC), trinucleotide based Moran autocorrelation (TMAC), dinucleotide based Geary autocorrelation (DGAC), trinucleotide based Geary autocorrelation (TGAC), and normalized Moreau-Broto autocorrelation (NMBAC) based on 38 properties. Likewise, the cross-correlation and auto cross-correlation module is further divided into two levels, dinucleotide-based cross-correlation (DCC) and trinucleotide-based cross-correlation (TCC), and dinucleotide-based auto-cross correlation (DACC) and trinucleotide-based auto cross-correlation (TACC). Finally, the pseudo correlation module is further divided into four sub-modules, parallel correlation pseudo dinucleotide composition (PC_PDNC), parallel correlation pseudo trinucleotide composition (PC_PTNC), serial correlation pseudo-dinucleotide composition (SC_PDNC), and serial correlation pseudo-trinucleotide composition (SC_PTNC). For DNA sequences, all the correlation-based features are generated, whereas, for RNA sequences, features incorporating only dinucleotide-based properties are calculated by Nfeature. Features based on dinucleotide properties include 38 properties, and trinucleotide properties-based features incorporate 12 properties. Our correlation-based module can calculate a total of 3,526 and 2,959 descriptors of DNA and RNA sequence (with default parameters).

#### Binary profile-based features

This module covers binary profile-based features of a given DNA/ RNA sequence. Binary profile-based features are important to motif predictions, factors/enhancers binding sites, etc. Using these modules, users can compute binary equivalents of each nucleotide in a given sequence. Generally, overlapping windows are used to create all possible patterns of fixed length for a given sequence. Then each pattern is converted to a binary profile (1 is used for presence and 0 is used to depict absence) as a numerical representation of each profile. Composition and correlation-based features are adequate to explain the function of the nucleic acids as a whole but fails to capture the residue level information. In order to predict the function of nucleic acids at the residue level, such as protein or biomolecule interacting nucleic acids, it is essential to represent the nucleotides in such a way or feature that apprehends the order and position of the nucleotides. Binary profiles or one-hot encoding meet the requirements mentioned above, where each or set of nucleotides designated by the vectors consist of ones and zeroes. In Nfeature, binary profiles are broadly categorized into two categories, such as property-based and nucleotide-based binary profiles. In DNA, property-based binary profiles are further divided into dinucleotide and tri-nucleotide property based binary profiles. In contrast, for RNA, only dinucleotide properties based binary profile is calculated. Whereas mono -, di- and tri-nucleotide based binary profiles are calculated for DNA and RNA under nucleotide-based binary profiles.

#### Nucleotide-Based Features

In this module, three sub-modules were involved for DNA and RNA, such as binary profile for mono-nucleotide (BPM), dinucleotide (BPD), and trinucleotide (BPT). In BPM, each nucleotide is represented by the vector of length four; for instance, A is defined as (1,0,0,0); G is designated as (0,1,0,0); C is indicated as (0,0,1,0), and T in case of DNA and U in case of RNA, is referred as (0,0,0,1). Similarly, for BPD, each dinucleotide is represented by the vector of size 16 with one element as 1, and the rest are zeros, such as dinucleotide AA is indicated as (1,0,0,0,0,0,0,0,0,0,0,0,0,0,0,0). Likewise, in BPT, each trinucleotide is represented by the vector of size 64. This module can compute 840 descriptors of a single DNA/RNA sequence with length of at least 10 residues.

#### Property-Based Features

This module is one of the novel modules of Nfeature, representing the binary profile of the sequence based on dinucleotide physicochemical properties (BP_DP) of DNA and RNA and also trinucleotide physicochemical properties (BP_TP) of DNA. 38 dinucleotide and 12 trinucleotide properties are involved in Nfeature. Each sequence in BP_DP is expressed by the vector of length 16*(N-1)*p, where N is the length of the sequence, and p is the number of selected features between 1 to 38. Likewise, the sequence in BP_TP is signified by the vector of length 64*(N-2)*p, where N is the length of the sequence, and p is the number of selected features between 1 to 12. The property-based module of Nfeature can compute highest number of features i.e., 11,616 of a single DNA sequence with length of at least 10 residues. In case of RNA based features, it can compute 6798 features of a single sequence with a length of 10 residue sequence.

#### Web Implementation and Standalone Package

In order to facilitate scientific community, we have developed a user-friendly webserver named as “Nfeature”. Webserver was integrated using Apache software on Linux/Ubuntu operating system. All the web pages have been developed with the help of HTML, CSS3 and PHP5. It is compatible with number of devices such as smart phone, laptop, Desktop, iPad. The submit page of server permit the user to submit nucleotide sequences (DNA/RNA) in FASTA format. The result page of web server allows the user to download the output in csv format. Figure 1 represents the description of all the modules of Nfeature tool (Figure 2). Additionally, the standalone package incorporates a readme file, description manual and separate codes for both DNA and RNA modules in respective directories.

**Figure 2:**
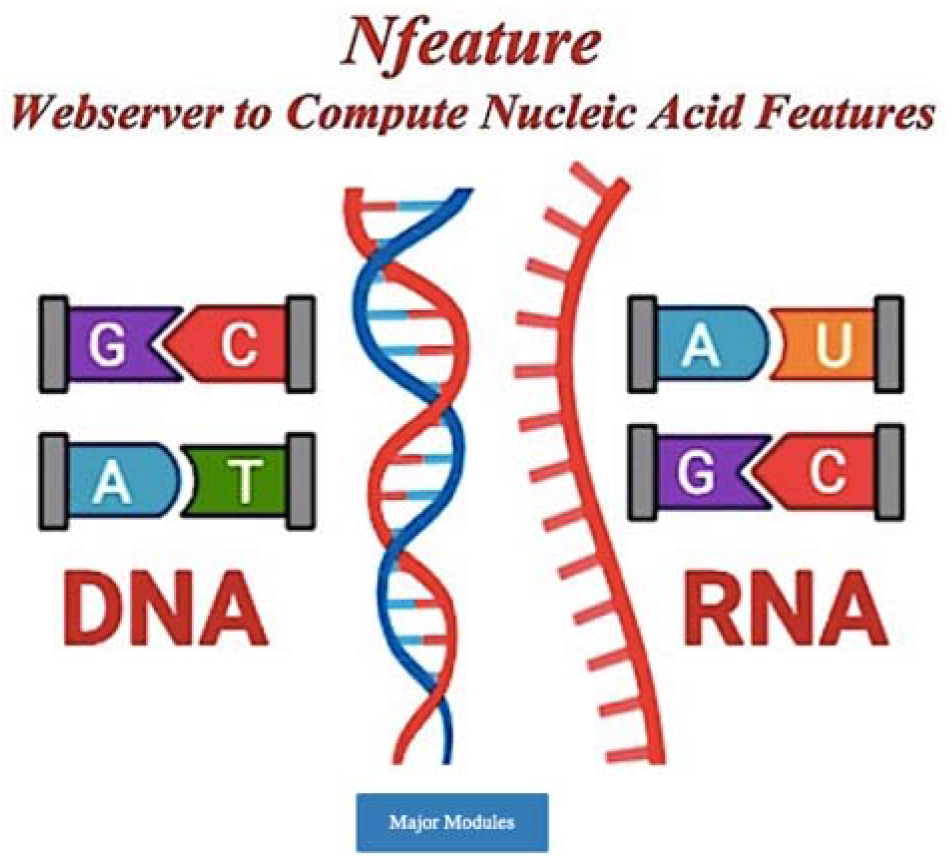
Web-interface of Nfeature to compute features of DNA/RNA sequences.

#### Comparison with other tools

In Table 1, we have compared Nfeature with the existing software/web servers based on the platform compatibility, package development and current running status. We showed that most of the software’s and packages are either not working or are not platform compatible with most of the frequently used operating systems. Nfeature and BioSeq-Analysis 2.0 are found to be available as a working web server and standalone package. Both of the tools were found to be compatible with widely used platforms such as Windows, Linux, and Mac OS.

**Table 1:** Comparison between the existing methods and Nfeature.

| Tools | Web Server | Standalone/Package | Platform Compatibility |
| --- | --- | --- | --- |
| Nfeature | ✓ | ✓ | Windows, Linux, and Mac OS |
| PseKNC | ✗ | ✗ | Not Working |
| **repDNA** | ✗ | ✓ | Windows, Linux, and Mac OS |
| **repRNA** | ✓ | ✗ | Windows, Mac OS |
| **BioTriangle** | ✓ | ✗ | Windows, Linux, and Mac OS |
| **BioSeq-Analysis2.0** | ✓ | ✓ | Windows, Linux, and Mac OS |
| **PyBioMed** | ✗ | ✓ | Windows, and Linux |

Software’s/web servers developed by various groups to compute various features based on input DNA/ RNA sequences, have their own limitations. Each tool/ web server aims to provide a unique set of features to serve user requirements. Nfeature is developed with an idea to add novel features to the existing features developed by other tools and also to provide conceivable feature generation techniques in a user-friendly fashion on a single platform. Nfeature integrates all the existing tools to calculate features for DNA and RNA sequences as shown in Table 2 and has added novel features.

**Table 2:**
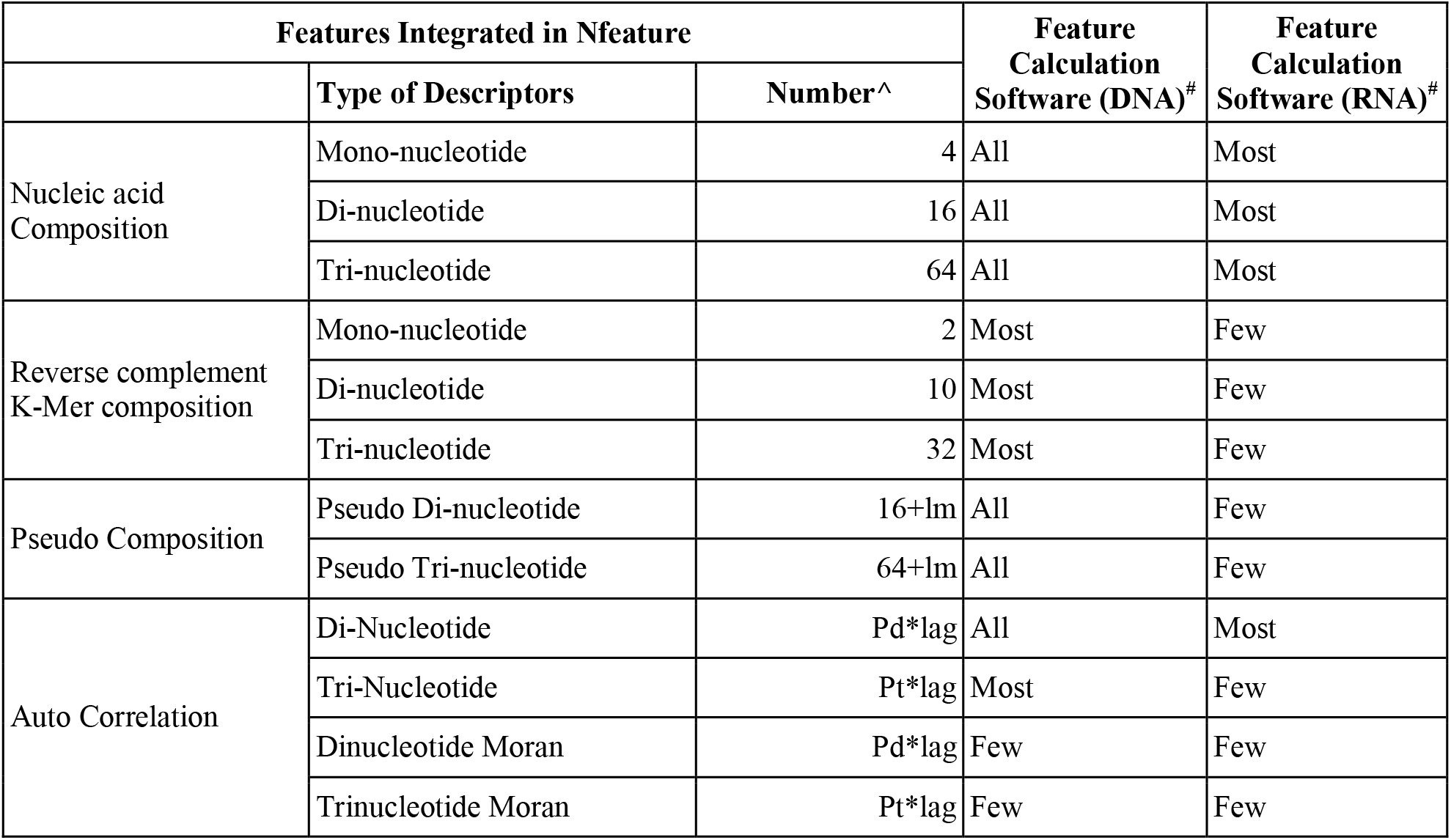
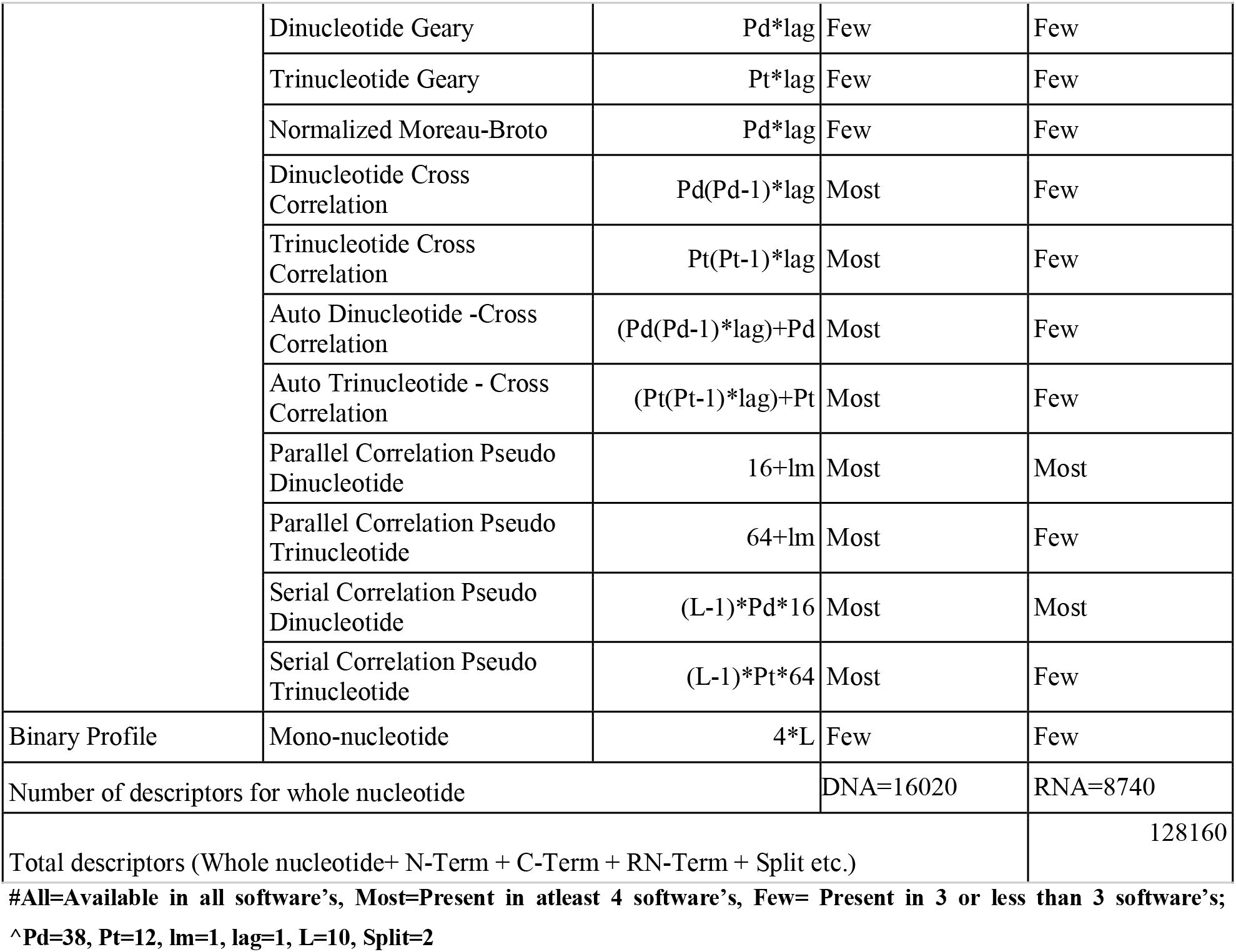
Comparison of features integrated in different platform/software. These descriptors are computed at nucleotide level, can be used to compute overall function/structure of a DNA/RNA sequence.

This platform provides new feature generation methods such as Nucleotide Repeat Index, Distance Distribution, Sequence level Entropy, Nucleotide level Entropy and Binary profiles of inputs DNA/ RNA sequences as represented in Table 3.

**Table 3:**
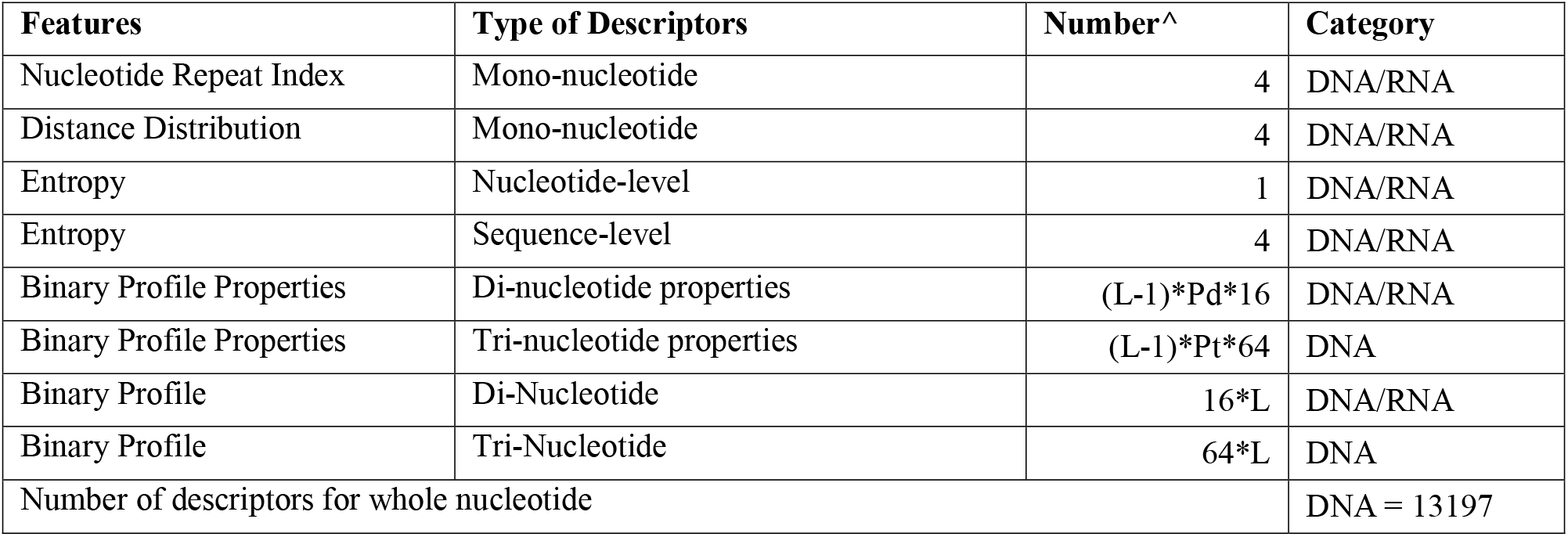
Novel features for DNA and RNA sequences incorporated in Nfeature Tool.

## Discussion and Conclusion

Due to advancement in technology in next generation sequencing, databases like NCBI (Coordinators, 2016), GenBank (Benson et al., 2013), EMBL (Stoesser et al., 2002), INSDC - DDBJ (Cochrane et al., 2016) are growing with exponential rate. In order to address numerous unsolved biological questions, there is an urgent need to develop computer-aided tools to annotate new sequences in above databases. In order to annotate any sequence, most important step is computation of numerical vector that represent characteristics of a sequence. In simple term computation of features or descriptors of a sequence is an important and essential step for computing function or structure of a sequence. In the past various package and web-based platform has been developed to compute wide range of features of proteins and nucleotide sequences. For example, pseudo K-tuple nucleotide composition (PseKNC), method used to generate composition and few correlation-based features from DNA/RNA sequences (Chen et al., 2014). RepDNA and RepRNA have been developed for calculating various features for DNA and RNA sequences respectively (Liu et al., 2015). BioTriangle is a web server that generate features for chemicals, proteins and nucleotide sequences and their interactions (Liu et al., 2016). They have reported to calculate 14 type features from DNA/RNA sequences.

PyBioMed also allows to generate features for chemicals, proteins nucleotides (Dong et al., 2016). This package generates compositions, autocorrelation and pseudo nucleic acid composition-based feature vectors. BioSeq-Analysis (Liu, 2019), which is recently updated to a new version, known as BioSeq-Analysis 2.0 (Liu et al., 2019) of the same was published including 26 features at the residue level and 90 features at the sequence level. This web server also develops the prediction models based on the feature generation technique, but the web server is very complex to use and not much user friendly. In addition, it doesn’t compute entropy-based features, Nucleotide repeat Index and Distance distribution of DNA/RNA sequences, unlike Nfeature. On the other hand, Nfeature integrates all existing features available for DNA/RNA sequences (Table 1) and adds eight novel features for better understanding of the sequence insights as represented in Table 2.

In summary, a number of tools have been developed to compute various features based on DNA/RNA sequences. These tools provide a unique set of features to cater different user requirements. Aim of developing Nfeature was to complement existing tools and to provide all possible feature generation techniques at a single platform in a user-friendly mode. Also, in order to overcome the limitations of the existing tools we have integrated all the existing tools features for DNA and RNA sequences with few new features in Nfeature platform. Overall, it is a comprehensive, easy-to-use web server/standalone package that allow users to calculate various features.

## Funding Source

The current work has not received any specific grant from any funding agencies.

## Conflict of Interest

The authors declare no competing financial and non-financial interests.

## Author contribution

MM and SP wrote all the scripts. AD, SP, RT, AA, and SJ developed the web interface. RT and AA prepared the manual. SJ and AD prepared the first draft of manuscript. MM, SP AD, SJ, RT, AA, and GPSR prepared the final version of manuscript. GPSR conceived the idea and coordinated the entire project.

## Acknowledgements

Authors are thankful to the Department of Computational Biology, IIIT-Delhi for infrastructure, Department of Biotechnology (DBT), Department of Science and Technology (DST-INSPIRE) and Council of Scientific and Industrial Research (CSIR), Govt. of India for financial support and fellowships.

## Notes

### Competing Interest Statement

The authors have declared no competing interest.

